# Water deficit response in nodulated soybean roots: a comprehensive transcriptome and translatome network analysis

**DOI:** 10.1101/2024.03.29.587361

**Authors:** María Martha Sainz, Carla V Filippi, Guillermo Eastman, Mariana Sotelo-Silveira, Sofía Zardo, Mauro Martínez-Moré, José Sotelo-Silveira, Omar Borsani

## Abstract

Soybean establishes a mutualistic interaction with nitrogen-fixing rhizobacteria, acquiring most of its nitrogen requirements through symbiotic nitrogen fixation. This crop is susceptible to water deficit; evidence suggests that its nodulation status—whether it is nodulated or not— can influence how it responds to water deficit. The translational control step of gene expression has proven relevant in plants subjected to water deficit. Here, we analyzed soybean roots’ differential responses to water deficit at transcriptional, translational, and mixed (transcriptional + translational) levels. Thus, the transcriptome and translatome of four combined-treated soybean roots were analyzed. We found hormone metabolism-related genes among the differentially expressed genes (DEGs) at the translatome level in nodulated and water-restricted plants. Also, weighted gene co-expression network analysis followed by differential expression analysis identified gene modules associated with nodulation and water deficit conditions. Protein-protein interaction network analysis was performed for subsets of mixed DEGs of the modules associated with the plant responses to nodulation, water deficit, or their combination. Our research reveals that the stand-out processes and pathways in the before-mentioned plant responses partially differ; terms related to glutathione metabolism and hormone signal transduction (2C protein phosphatases) were associated with the response to water deficit, terms related to transmembrane transport, response to abscisic acid, pigment metabolic process were associated with the response to nodulation plus water deficit. Still, two processes were common: galactose metabolism and branched-chain amino acid catabolism. A comprehensive analysis of these processes could lead to identifying new sources of tolerance to drought in soybean.

## 1. Introduction

Leguminous plants such as soybean (*Glycine max* (L.) Merr.) can establish a root nodule endosymbiosis with nitrogen-fixing soil rhizobacteria. This mutualistic interaction induces significant metabolic and nutritional changes in the plant (Oldroyd et al., 2011; Concha and Doerner, 2020). The symbiotic nodule, the newly developed root organ due to the symbiotic pathway, is where nitrogen fixation and assimilation occur—a process known as symbiotic nitrogen fixation (SNF). The main products of SNF, ureid or amide compounds in determinate or indeterminate nodules, respectively, are exported to the rest of the plant, which in turn provides the rhizobia with photoassimilates (Van Heerden et al., 2007). Since developing and maintaining nodules is resource-consuming, the plant exerts tight control over the nodulation process under favorable and sub-optimal growing conditions (Ferguson et al., 2019).

Evidence suggests that the nodulation condition of a legume (i.e., nodulated or non-nodulated) can affect its response strategies to water deficit (Antolín et al., 1995; Borsani et al., 1999; Lodeiro et al., 2000; Staudinger et al., 2016; Liu et al., 2022); however, it is uncertain which molecular mechanisms are responsible for the differential response. The regulation of gene expression, which can be achieved at the transcriptional and/or post-transcriptional levels (including translational and post-translational events), could explain the plant differential response strategies previously mentioned. In particular, translational control has proven relevant in plants subjected to stressful conditions such as nutrient scarcity (Bazin et al., 2017) or different situations of biotic and abiotic stresses like water deficit (Kawaguchi et al., 2004; Lei et al., 2015). Plants benefit from this step of gene expression regulation, which does not require de novo messenger RNA (mRNA) synthesis but rather refers to the efficiency with which mRNAs already present in cells are translated since it allows them to respond rapidly, thus conferring flexibility and adaptability (Lee and Bailey-serres, 2019; Urquidi Camacho et al., 2020). At present, it is well established that the poor or variable levels of correlation between the levels of transcripts and proteins found in different organisms is explained by the post-transcriptional steps of gene expression, mainly translation (Piccirillo et al., 2014; Lei et al., 2015; Becker et al., 2018; Traubenik et al., 2020). This way, the direct analysis of the subset of mRNAs that are being translated (the translatome) enables more accurate and complete measurement of cell gene expression than the one obtained when only the transcriptome (steady-state mRNA levels) is analyzed (Sablok et al., 2017). Still, the information gathered from a transcriptomic and a translatomic analysis is complementary, and their comparison allows, for instance, to distinguish between the different levels of regulation of each mRNA.

Despite how advantageous translational control is for plants subjected to stressful conditions, studies that involve a translatomic analysis of nodulated and water-restricted plants are scarce. As an interesting example, we can mention the work recently published by our group, in which we reported that some members of the thioredoxin and glutaredoxin systems were regulated mainly at the translational level in the roots of nodulated soybean plants subjected to water-deficit stress (Sainz et al., 2022a), thus resignifying the relevance of these enzymes for having a unique role in nodulated plants subjected to water restriction.

The plant roots are the organs first exposed to water deficit and are thus responsible for sensing water shortage, a main constraint for crop production, and transmitting stress signals to the rest of the plant. This causes developmental, physiological, and metabolic changes in adaptation to the water deficit, leading to the acquisition of resistance at the whole-plant level (Takahashi et al., 2020; Kang et al., 2022). The responses of plant roots, primarily those of crop plants, have therefore been the subject of research over the past few decades, and the selection of plants with root traits that improve productivity under drought is of great relevance for geneticists and breeders (Comas et al., 2013).

This study aimed to analyze the differential responses of roots of nodulated soybean plants to water deficit by evaluating gene expression regulation at the transcriptional, translational, or mixed (transcriptional + translational) levels. Thus, the transcriptome (total RNA fraction) and translatome (polysome-associated RNA fraction) of four combined-treated soybean roots (including the nodulation and water deficit conditions) were analyzed. Genes encoding various enzymes involved in hormone metabolism were found among the differentially expressed genes (DEGs) mainly regulated at the translatome level in nodulated and water-restricted plants. Further, we identified gene modules associated with the nodulation and/or water deficit conditions of soybean plant roots through a weighted gene co-expression network analysis (WGCNA) followed by a differential expression analysis (Sánchez-Baizán et al., 2022). Gene modules more representative of the DEGs were subjected to enrichment analysis, and selected modules were further analyzed. Protein-protein interaction network analysis was performed for subsets of DEGs of the modules more associated with the plant responses to nodulation, water deficit, or the combination of nodulation + water deficit, highlighting the stand-out biological processes and metabolic pathways in the before-mentioned plant responses.

## 2. Results

### 2.1 Scope of the experimental design and data quality

Our experimental design comprised soybean plants subjected to four treatments as a result of combining two nodulation conditions, i.e., nodulated (N) and non-nodulated (NN) plants, with two hydric conditions, i.e., water-restricted (WR) and well-watered (WW) plants. While this study aimed to analyze the responses of nodulated and water-restricted plants, this comprehensive experimental design allowed us to perform several comparisons between the before-mentioned treatments to investigate the distinctive responses of plants to the different nodulation and hydric conditions.

In this study, we analyzed the following four comparisons: *i*) N+WR *vs*. N+WW; *ii*) N+WR *vs*. NN+WR; *iii*) NN+WR *vs.* NN+WW; *iv*) N+WW *vs*. NN+WW (Figure 1). The N+WR *vs*. N+WW (*i*) comparison exhibits the response of nodulated plants to water deficit; meanwhile, the *ii* comparison (N+WR *vs*. NN+WR) shows the particular response to water deficit of nodulated plants with respect to non-nodulated plants. The NN+WR *vs.* NN+WW (*iii*) comparison evidence the response of non-nodulated plants to water restriction, and the *iv* comparison (N+WW *vs*. NN+WW) shows the plant responses due to the nodulation process without involving water restriction.

**Figure 1.**
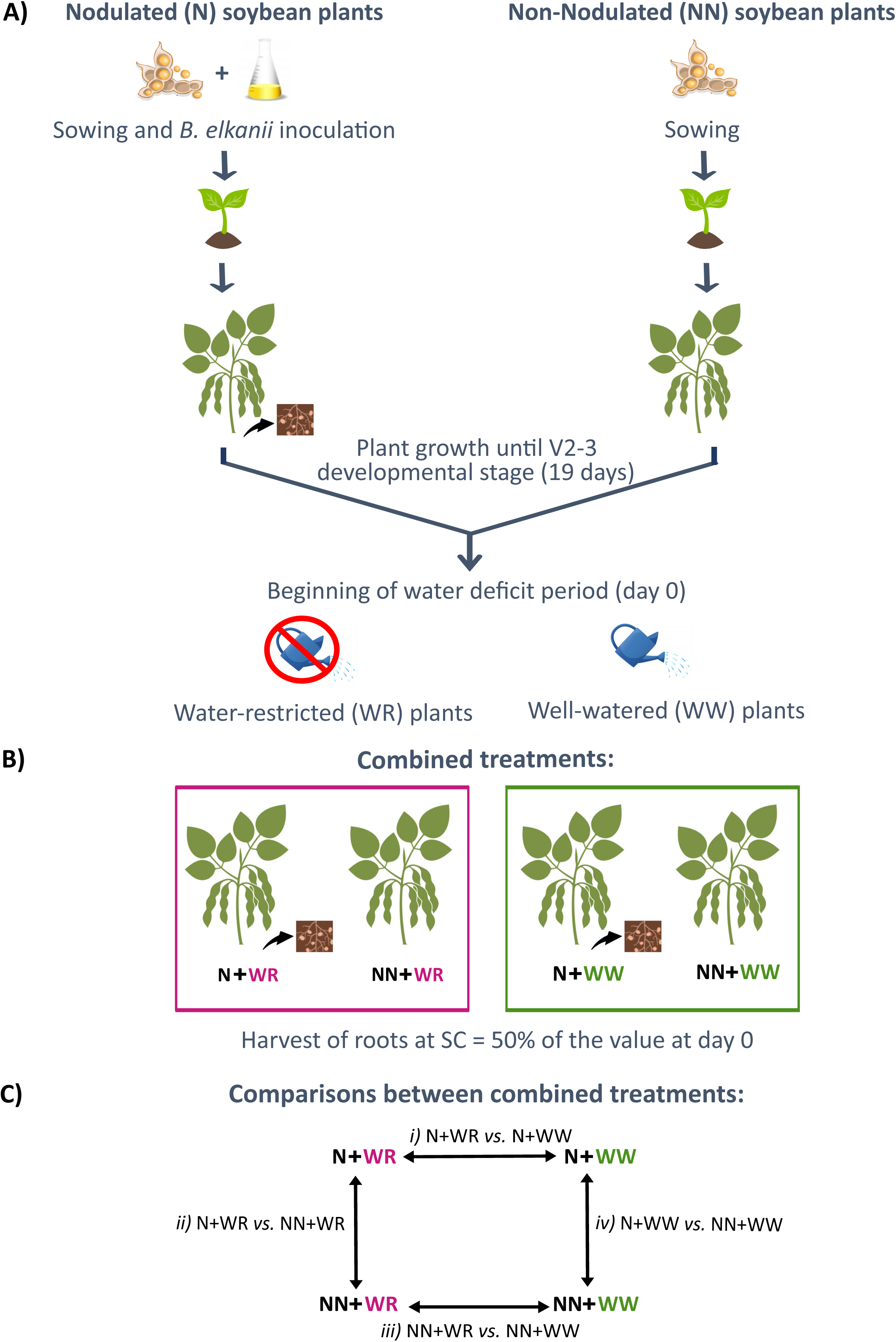
Schematic overview of the methodology followed for the obtention of the combined treated soybean roots and the comparisons between the combined treatments. **(A)** The scheme gives an overview of the steps followed from plant growth, inoculation (only for nodulated plants), and the establishment of the water deficit stress (only for water-restricted plants). **(B)** As a result, soybean plants subjected to four combined treatments were obtained: nodulated and water-restricted plants (N+WR), non-nodulated and water-restricted plants (NN+WR), nodulated and well-watered plants (N+WW), non-nodulated and well-watered plants (NN+WW). **(C)** Four comparisons between combined treatments were analyzed: *i*) N+WR *vs*. N+WW; *ii*) N+WR *vs*. NN+WR; *iii*) NN+WR *vs.* NN+WW; *iv*) N+WW *vs*. NN+WW. N: nodulated; NN: non-nodulated; WR: water-restricted; WW: well-watered: SC: stomatal conductance.

The four comparisons were studied at the transcriptome (total RNA fraction; TOTAL) and the Polysome-Associated RNA level (PAR fraction) to analyze the plant responses to the different nodulation and hydric conditions in a comparative manner and to highlight changes at the loading of mRNA in the translational apparatus versus total RNA levels. The latter allows to assess regulation at the transcriptome (TOTAL) and translatome (PAR) levels as well as mixed responses (TOTAL+PAR) involving those compartments upon the conditions compared.

Exploration of the RNA-seq data using distance matrix analysis (heatmap) and principal component analysis (PCA) showed that all the biological replicates clustered together (Figure S1, A and B). Also, the PCA evidenced that component 1 (PC1) explained the greatest proportion of the variance (36%) separating the samples by hydric condition (pink *vs.* green). Component 2 (PC2) separated the samples by the nodulation condition (circles *vs.* triangles). Thus, four well-defined groups were obtained, corresponding to the four combined treatments. As expected, each sample’s TOTAL and PAR RNA fractions were found to be very close to each other (Figure S1, B).

### 2.2 The metabolism of several hormones is regulated at the translational level in nodulated and water-restricted plants

The results of the exploratory analysis of the RNA-seq data suggested that the data obtained were suitable for further downstream analysis. In this line, to learn how the nodulation condition and the water-restriction condition affect the plant’s transcriptome and translatome, an initial differentially expressed gene (DEG) analysis was performed contrasting nodulated samples with respect to non-nodulated samples (including WR and WW plants) on one side (Figure 2, A and B), and water-restricted samples with respect to well-watered samples (including N and NN plants) on the other (Figure 2, C and D). Overall, water restriction significantly impacted gene expression at the transcriptome and translatome level more than nodulation. In the N *vs.* NN comparison, most DEGs were upregulated, while in the WR *vs.* WW comparison, more DEGs were downregulated at both TOTAL and PAR levels (Figure 2, A, B, C, D).

**Figure 2.**
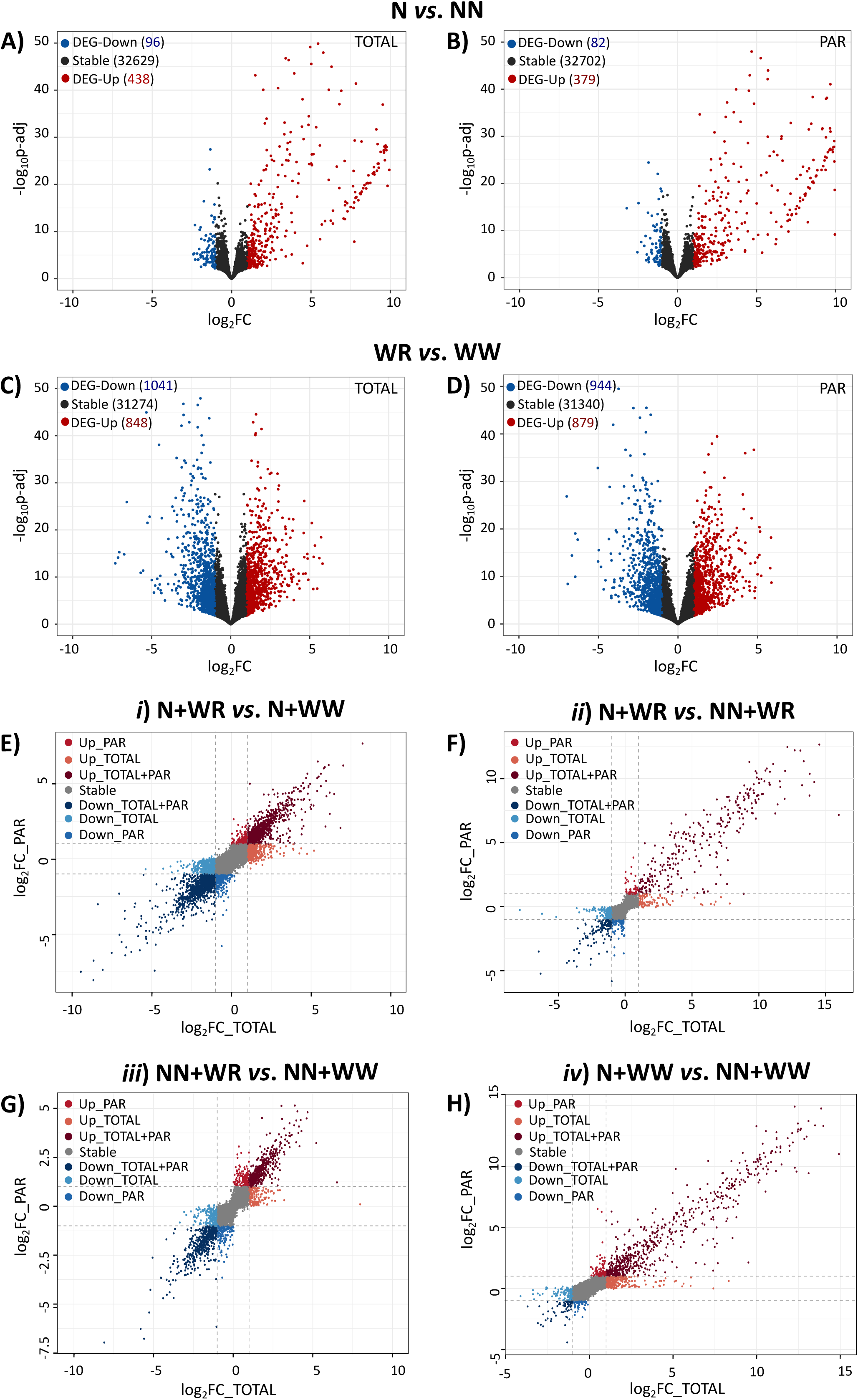
Transcriptional and translational responses of soybean roots to nodulation and water restriction. Volcano plots showing stable genes, down- and up-regulated DEGs in the contrast between nodulated *vs.* non-nodulated plants at the TOTAL **(A)** and PAR **(B)** levels. Volcano plots showing stable genes, down- and up-regulated DEGs in the contrast between water-restricted *vs.* well-watered plants at the TOTAL **(C)** and PAR **(D)** levels. Scatterplots showing the FC in the TOTAL and PAR samples in the comparisons between the combined treatments analyzed: *i)* N+WR *vs.* N+WW **(E)**; *ii)* N+WR *vs.* NN+WR **(F)**; *iii)* NN+WR *vs.* NN+WW **(G)**; *iv)* N+WW *vs.* NN+WW **(H)**. Each dot represents the log_2_ of the FC in TOTAL and PAR samples. DEGs are shown as colored dots.

To further identify DEGs with translational regulation in each of the four comparisons between combined treatments analyzed, scatter plots showing the fold change (FC) in the corresponding TOTAL and PAR samples were done (Figure 2, E, F, G, H). Interestingly, the set of DEGs with the highest FC regulated mainly at the PAR level in nodulated and water-restricted plants (comparisons *i* and *ii*; Figure 2, E, F; Table S2) includes many genes related to the metabolism of the hormones abscisic acid (ABA), ethylene, auxin and cytokinin such as those encoding the zeaxanthin epoxidase, non-race-specific disease resistance/hairpin-induced (NDR1/HIN1)-like protein 6, 1-aminocyclopropoane-1-carboxylate (ACC) synthase 3, indole-3-acetic acid (IAA)-amido synthetase GH3.10, indole-3-acetate O-methyltransferase 1 (IAMT1), and adenylate isopentenyltransferase 5 (IPT5). Given the key role of these hormones in the plant response mechanisms to the processes under investigation (nodulation and water restriction), the fact that these genes were differentially expressed at the translational level in N+WR plants is highly pertinent. Genes encoding amino acids and sucrose transporters, cell wall proteins and enzymes related to its modification, plant receptor-like kinases (RLKs), protein phosphatases 2C, heat shock proteins (HSPs), and transcription factors (WRKY 72A, MADS-box 23, bHLH18, WUSCHEL-related homeobox 4) were also included among those DEGs mainly regulated at the PAR level in N+WR plants (Figure 2, E, F; Table S2). Examples of DEGs solely regulated at the translational level in comparison *iii* (the one not including nodulated plants) comprised those encoding late embryogenesis abundant (LEA) proteins, heat shock proteins (HSP), inositol phosphate synthase—a rate-limiting enzyme in myo-inositol biosynthesis—transcription factors, and ABA and auxin metabolic enzymes (Figure 2, G; Table S2). Comparison *iv* (which only involved the nodulation condition) had among its DEGs mainly regulated at the PAR level those coding for enzymes related to cell wall reconstruction or modification, cytokinins and phenylpropanoid biosynthesis, and amino acids transporters, among others (Figure 2, H; Table S2).

Moreover, the presence of some genes in nodulated and water restricted plants (particularly highlighted in contrast *ii*) with a notable transcriptional upregulation (Up-TOTAL; the ones with the higher FC at the TOTAL level) is another intriguing aspect depicted in Figure 2 (Figure 2, F; Table S2). Strikingly, despite their transcriptional upregulation, these transcripts seem to be hindered from associating with polysomes and undergoing active translation. This fact can suggest a potential sequestration of these transcripts within biomolecular condensates, such as P bodies or stress granules (Parker et al., 2022), offering a convincing avenue for additional research.

### 2.3 Weighted gene co-expression network analysis identified 25 gene modules with coordinated changes in the combined-treated soybean roots

WGCNA was performed using the 24 samples comprising the four combined-treated roots (in triplicates) and each sample’s TOTAL and PAR fractions. The analysis identified 25 co-expression modules, as shown in Figure 3A, using 26,793 genes, which were the ones left after genes with low coefficients of variation and/or low counts were removed (see 4.4.3; Table S3). These 25 distinct expression modules contained between 27 and 6276 genes and were further merged into seven groups based on their eigengene expression patterns (Figure 3A). As can be seen, the distinct groups were differentially detected in the different samples and RNA fractions. It is worth mentioning that the first group (ME5, ME1, ME14, ME13, ME16, ME22) was mainly detected in polysome-associated RNA fraction across all treatments, with the highest detection in non-nodulated plants without water restriction (NN+WW treatment). The second group (ME11, ME9, ME12) exhibited detection in both RNA fractions (TOTAL and PAR) of the N+WW treatment, specifically associated with nodulation. Group three (ME18, ME7, ME24, ME20) included genes preferentially detected in well-watered conditions (NN+WW and N+WW treatments). Notably, the highest and lowest transcript detection were observed in NN+WW and N+WR treatments, respectively, highlighting the specificity of these genes to the treatment combination involving the most contrasting conditions. Group six (ME21, ME6, ME3, ME10, ME23) exhibited higher detection in N+WR and NN+WR treatments compared to the other two well-watered combined treatments. This is graphically presented in Figure 3A.

**Figure 3.**
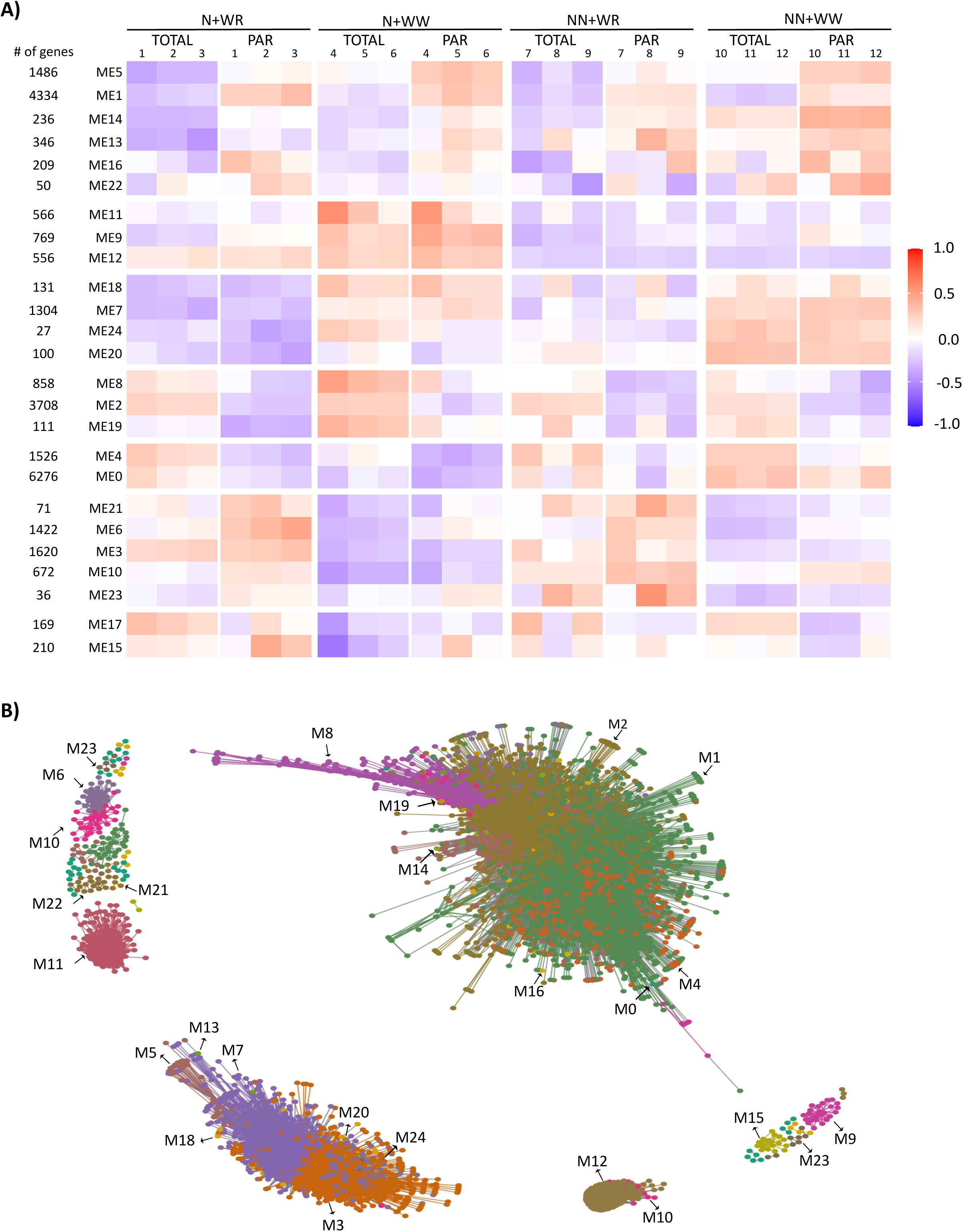
Gene co-expression network of soybean plants subjected to four combined treatments comprising nodulation and water restriction conditions and total RNA and polysome-associated mRNA samples. **(A)** Heatmap of module eigengene (ME) expression profiles in the indicated combined treatments and RNA fraction (TOTAL, PAR). The three biological replicates of each treatment and the number of genes in each module are also shown. **B)** Projection of expression modules (M) of the weighted gene co-expression network where 24 out of 25 modules are visualized; M17 is not visualized due to a low edge adjacency value. The network contained 8,944 nodes and 1,378,580 edges disposed as a central core and several smaller sub-networks disconnected from the central core.

The topological representation of the expression modules of the WGCNA network was projected in different colors using the R package network (Butts, 2008) (Figure 3B). All modules, except for M17, showed edge adjacency values higher than the threshold (0.15) and thus are present on the network. This way, the whole network contained 8,944 nodes and 1,378,580 edges, disposed of as a central core, and several smaller sub-networks disconnected from the central core (Figure 3B).

### 2.3 DEG analysis identified eight co-expressed modules as the more relevant ones in the plant responses to nodulation and/or water deficit

To explore further the plant responses to the different experimental conditions, the DEGs and their status (i.e., up- or down-regulated) found in the four comparisons between the combined treatments (Figure 1C) were highlighted on the WGCNA network (Figure 4). Moreover, the number of genes present in the three regulation levels analyzed (TOTAL, PAR, and TOTAL+PAR) was further discriminated in Table S4. Comparison *i*, which exhibited the response of nodulated plants to water deficit, had the greatest number of DEGs that localized within specific modules. M3, M5, M6, M7, M9, M10, M11, M12, M18, and M19 were the most representative modules in this case and included a similar number of up- and down-regulated DEGs (Figure 4A; Table S4). The DEGs obtained in comparison *ii*, which showed how plants change their response to water deficit when involved in a symbiotic interaction with rhizobia, belonged mainly to modules 3, 7, and 13 (Figure 4B). In this case, most DEGs (84%) were down-regulated (blue dots; Figure 4B; Table S4). Comparison *iii* (NN+WR *vs.* NN+WW) evidenced the response of soybean plants to water restriction; here, the DEGs were mainly localized within M5, M7, M12, and M23 and comprised a similar number of up- and down-regulated DEGs (Figure 4C; Table S4). The comparison that showed the plant responses due to the nodulation process without involving any water restriction, comparison *iv*, presented its DEGs principally in modules 9, 10, 11, and 12 (Figure 4D), with 72% of them being up-regulated (red dots; Figure 4D; Table S4). Notably, the most contrasting situation regarding the status of the DEGs was observed between comparison *i* and *iv* (Figure 4A and D, respectively), where DEGs within modules 9, 10, 11, and 12 exhibited opposite regulation; e.g., DEGs in M11 were up-regulated in comparison *iv* whereas down-regulated in comparison *i*. Hence, many DEGs up-regulated due to the nodulation process inverted their expression when the water deficit was imposed on the nodulated plants.

**Figure 4.**
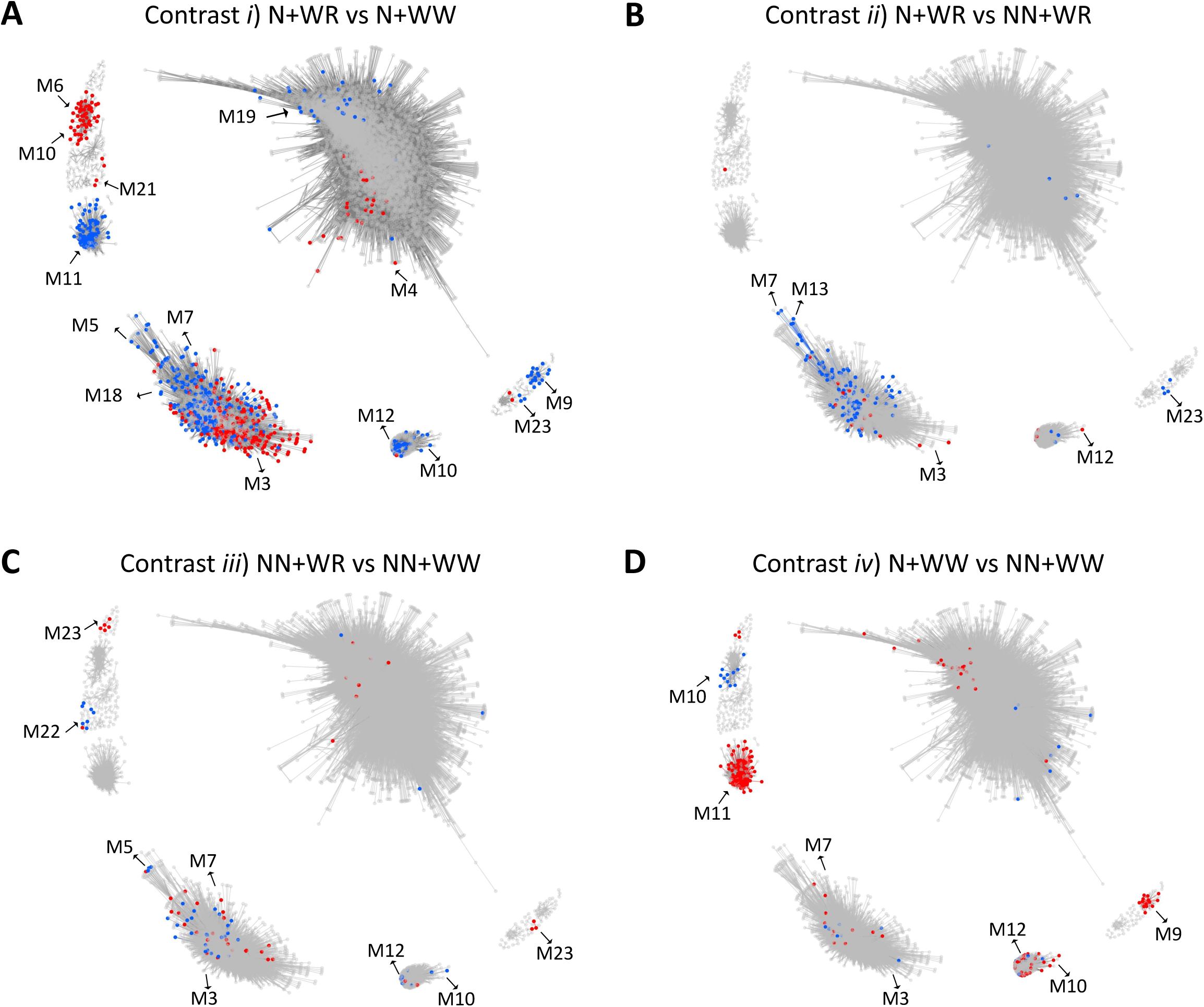
Identification of differentially expressed genes (DEGs) at TOTAL, PAR, and TOTAL+PAR regulation levels in the contrasts between combined treatments within the WGCNA network. **(A)** Contrast *i*) N+WR *vs.* N+WW. **(B)** Contrast *ii*) N+WR *vs.* NN+WR. **(C)** Contrast *iii*) NN+WR *vs.* NN+WW. **(D)** Contrast *iv*) N+WW *vs.* NN+WW. Red dots represent up-regulated DEGs. Blue dots represent down-regulated DEGs. The four combined treatments were nodulated and water-restricted plants (N+WR), nodulated and well-watered plants (N+WW), non-nodulated and water-restricted plants (NN+WR), non-nodulated and well-watered plants (NN+WW). M: module.

Based on the number of DEGs localized at each of the WGCNA modules (Table S4), the eight most representative ones (M3, M5, M6, M7, M9, M10, M11, and M12) were selected for further analysis (Table 1). Table 1 complements the data presented in Figure 4 by specifying the regulation at TOTAL, PAR, or TOTAL+PAR level for each comparison in which the DEG condition was achieved, at each status, and for each of the eight selected modules. Moreover, in Table 1, the number of the modules’ hub genes (10% most connected) is shown in parentheses since we wondered whether any of the DEGs corresponded to any of the hub genes in each modules, as highly connected genes are often more important for the functionality of networks that other nodes (van Dam et al., 2018). In fact, in three out of the four comparisons analyzed, some DEGs were also hub genes, with the N+WR *vs.* NN+WR comparison (*ii*) having the most hub genes regarding its DEGs (12%: 25 out of 214). Comparison *iii* (NN+WR *vs.* NN+WW) contained no hub genes among their DEGs. Furthermore, compared to the percentage found for the just mentioned comparisons *ii* (12%) and *iii* (0%), comparisons *i* (N+WR *vs*. N+WW) and *iv* (N+WW *vs*. NN+WW) showed an intermediate percentage of hub genes among their DEGs (4%: 70 out of 1636, and 5%: 21 out of 410, respectively). Our findings suggest that, whereas the nodulation condition affected the differential expression of some hub genes, the interaction between nodulation and water deficit (relative to non-nodulation) affected the differential expression of a higher proportion of hub genes (Table 1).

**Table 1.**
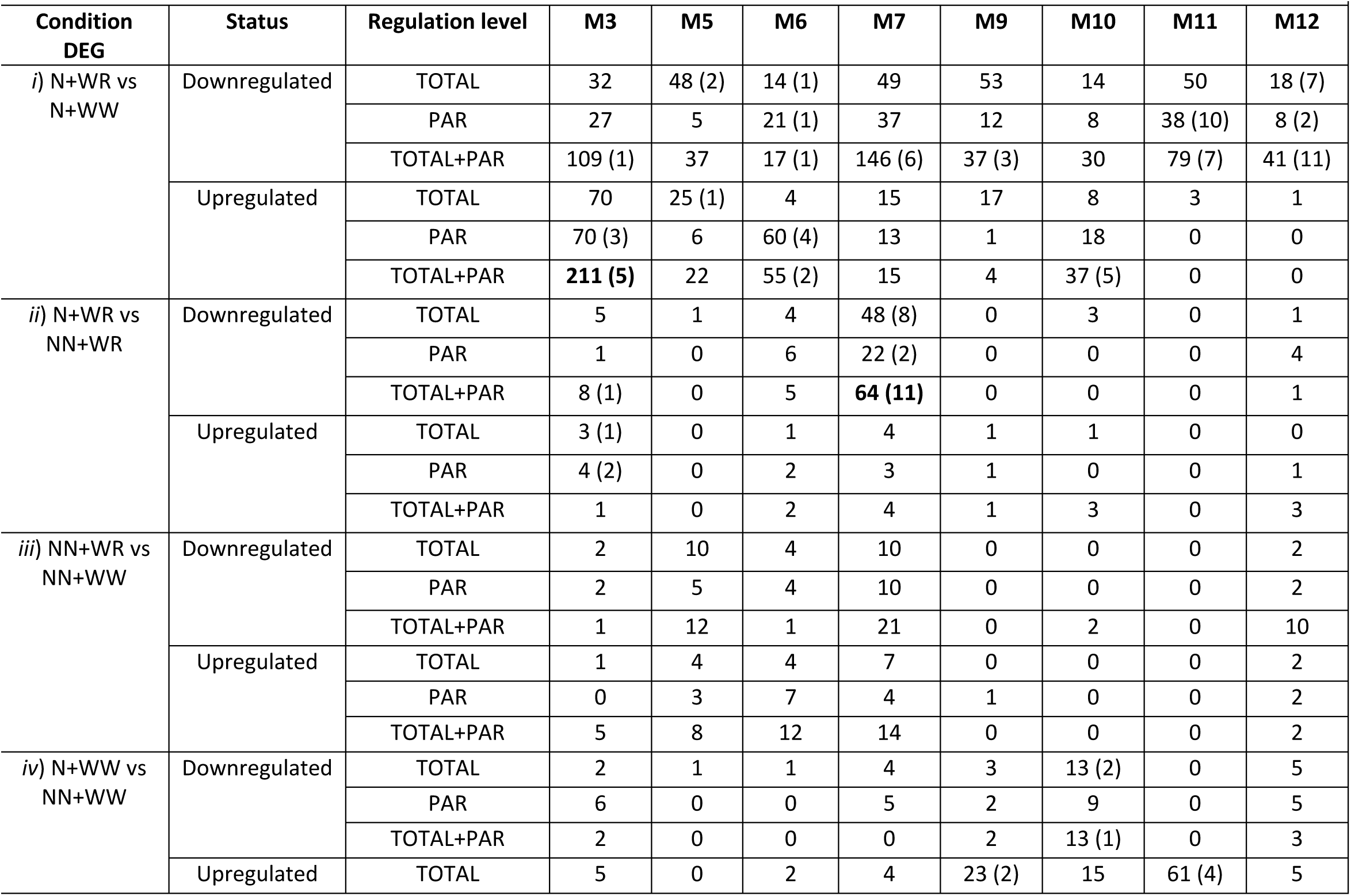
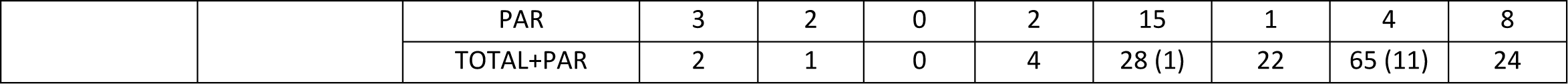
Weighted gene co-expression network analysis (WGCNA) modules (M) most representative of the differentially expressed genes (DEGs) obtained in the four (*i*, *ii*, *iii*, *iv*) comparisons analysed. M3, M5, M6, M7, M9, M10, M11 and M12 are depicted. The status (up- or down-regulated) and the regulation level (TOTAL, PAR, or TOTAL+PAR) are also shown. The number of the modules’ hub genes is presented in parentheses. The hub genes were defined as the 10% of genes with the highest connectivity.

Another interesting focus point regarding our WGCNA+DEG analysis is the common DEGs among two comparisons that are discriminated according to their expression modules. The common DEGs found between comparisons *i* and *iii* are genes related to the plant responses to the water deficit, independently of the nodulation condition (Figure 2C and D). Instead, the ones found in common between comparisons *ii* and *iv* are genes related to nodulation, regardless of the hydric condition (Figure 2A and B). In both cases, there were a significant number of DEGs at TOTAL+PAR regulation level in some modules (Table S5). Interestingly, these DEGs were specific to certain co-expression modules (M3, M5, M6, M7, M12) and were hub genes in some cases (Table 2). Moreover, the common DEGs between comparisons *i* and *iii* were mainly present in M3, M5, M6, and M7, whereas the common DEGs between comparisons *ii* and *iv* were almost entirely found in M12 and were almost entirely upregulated (Table 2). It is noteworthy that 27% (30 out of 113) of these upregulated DEGs were also hub genes of the module. In conclusion, most genes related to the plant responses to the water deficit were localized within four specific WGCNA expression modules (M3, M5, M6, M7), but, among these, mainly within M3 and M7. Also, almost all the genes related to nodulation were specifically localized in M12.

**Table 2.**
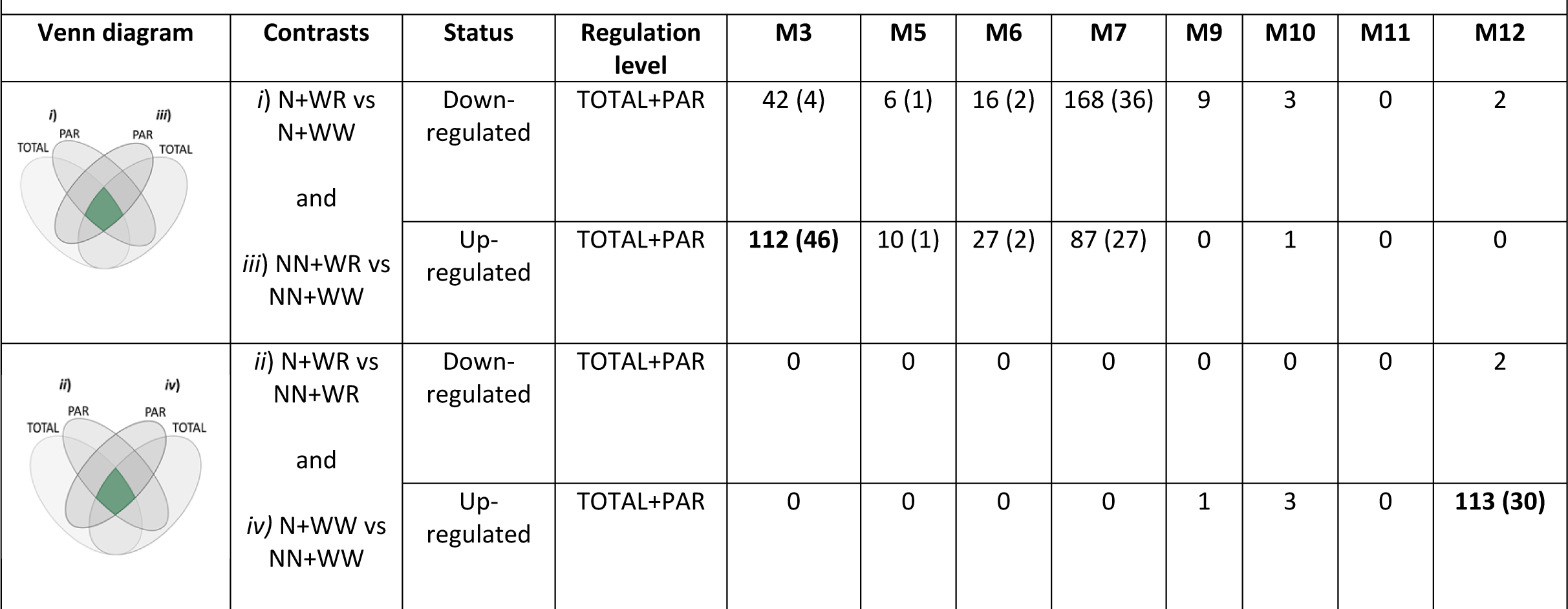
Differentially expressed genes (DEGs) common to contrasts *i*) and *iii*) and contrasts *ii*) and *iv*) at TOTAL+PAR level depicted according to their localization at the WGCNA modules shown in Table 1 (M3, M5, M6, M7, M9, M10, M11, and M12). The Venn diagrams are shown for clarity; the green area within represents the intersection of common DEGs at the TOTAL+PAR level between the respective contrasts. Also, the status (up- or down-regulated) is shown. The number of the modules’ hub genes is presented in parentheses. The hub genes were defined as the 10% of genes with the highest connectivity.

M3 comprised 67 hub genes within the network; 60 were DEGs (Table S3). Among them, 50 were common between comparisons *i* and *iii* at TOTAL+PAR level, and most (46) were up-regulated (Table 2). M5 and M6 each had 21 hub genes; five DEGs in M5 and 13 in M6 (Table S3). In M7, 75 out of 78 hub genes in M7 were DEGs, with 63 common in comparisons *i* and *iii* at TOTAL+PAR level (Table 2). M9 contained three hub genes coding histone H3.2 (Table S3). All five hub genes in M10 were up-regulated in N+WR plants (regarding N+WW plants, comparison *i*) at TOTAL+PAR level. Also, three were down-regulated in comparison *ii* (Table S3; Table 1). Regarding M11, the 18 hub genes were DEGs (Table S3), with down-regulation in comparison *i* at PAR and TOTAL+PAR levels and up-regulation in comparison *iv* at TOTAL and TOTAL+PAR levels (Table 1). Notably, 33% (6 out of 18) of M11 hub genes were transcription factors (TFs) (Table S3). All 30 M12’ hub genes were up-regulated DEGs common between comparisons *ii* and *iv* at TOTAL+PAR level (Table 2), and 20 were down-regulated DEGs in comparison *i* at TOTAL, PAR, and TOTAL+PAR levels (Table 1).

### 2.4 Characterization of biological processes and pathways in the selected WGCNA modules

Subsequently, Gene Ontology (GO) functional enrichment analysis (biological process; BP) was used to identify and rank overrepresented functional categories (van Dam et al., 2018) of the previously selected modules (M3, M5, M6, M7, M9, M10, M11, and M12) (Figure S2). M3, M7, M11, and M12 were chosen for further interpretation based on the DEGs characteristics (shown in Table 1 and Table 2) and on our interest in the BP and pathways in which they are enriched (Figure 5). In addition to the GO_PB shown in Figure S2, this figure depicts the Kyoto Encyclopedia of Genes and Genomes (KEGG) pathway enrichment analysis and the modules network with the hub genes highlighted in the central part. The different topologies of the networks, together with the centrality of the hub genes (big dots), can be appreciated. Notably, the nodes (small dots) in M12’ network displayed a sharp peripheral distribution, while the networks of the other modules had a more uniform node distribution (Figure 5).

**Figure 5.**
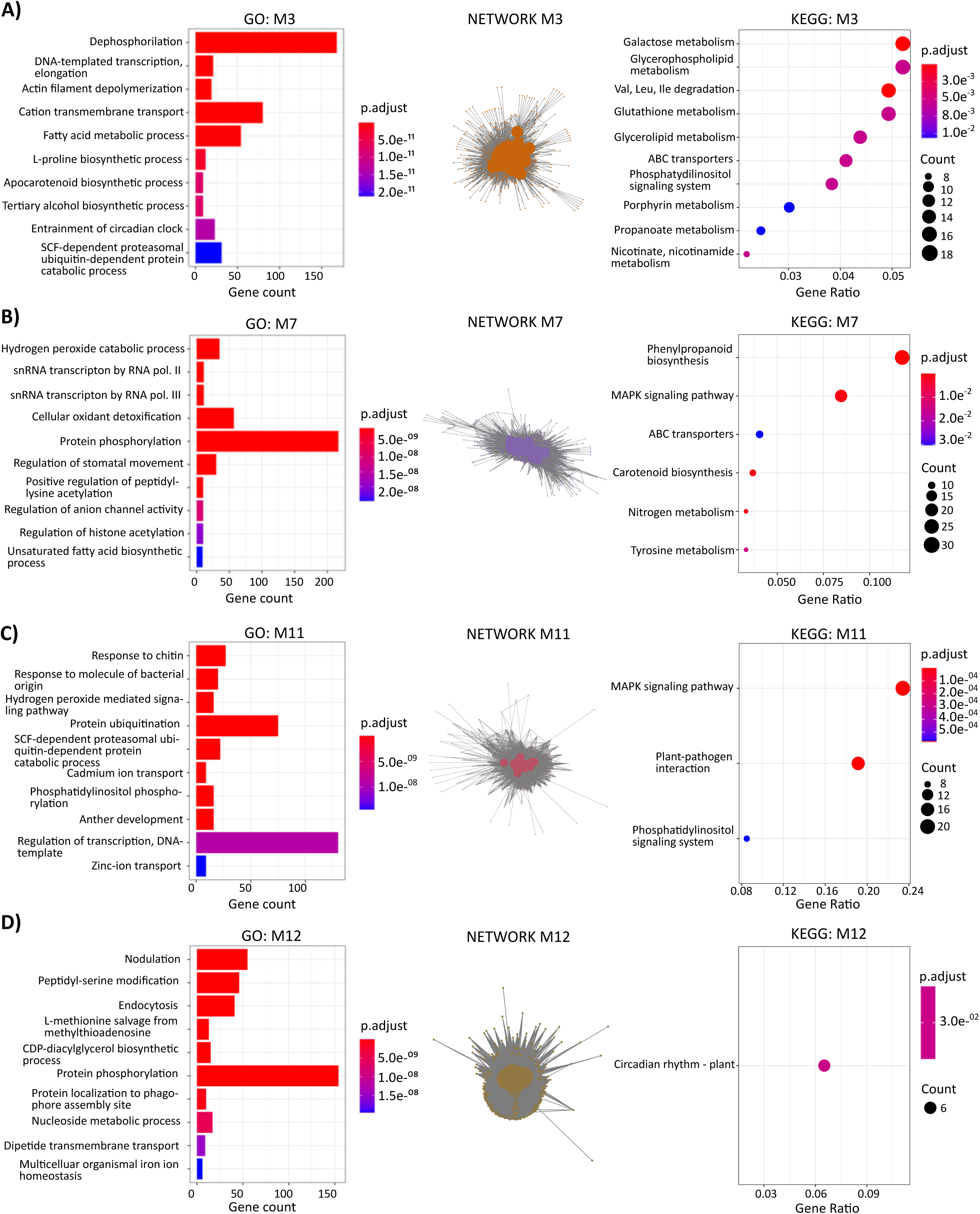
Functional and pathway enrichment analysis and network representation of co-expression modules M3 (A), M7 (B), M11 (C) and M12 (D). The top ten Gene Ontology Biological Process (GO-BP) terms are shown for each module (y-axis, left chart). The nodes of the networks are highlighted with the colour of the corresponding module (according to Figure 3B), and hub genes are the ones with the bigger dots. The modules’ KEGG-enriched pathways are depicted in the right chart. The hub genes were defined as the 10% of genes with the highest connectivity.

Module 3, closely related to the response of nodulated plants to water deficit, was strongly associated with dephosphorylation (GO:0016311), cation transmembrane transport (GO: 0098655), and fatty acid metabolism (GO:0006631) processes (Figure 5A). The KEGG pathways with higher gene counts included galactose and glycerophospholipid metabolisms, degradation of branched-chain amino acids (BCAAs), and glutathione (GSH) metabolism (Figure 5A). M7 was enriched in protein phosphorylation (GO:0006351), cellular oxidant detoxification (GO:0098869), and hydrogen peroxide catabolism (GO:0042744) processes, the first of which had over 200 gene counts (Figure 5B). Moreover, KEGG enrichment highlighted phenylpropanoid biosynthesis and the MAPK signaling pathway (Figure 5B). The two GO-BPs with the greatest gene counts in M11 were regulation of transcription (GO:0006351) and protein ubiquitination (GO:0016567) (Figure 5C). KEGG analysis, in this case, showed enrichment in the MAPK signaling pathway, plant-pathogen interaction, and phosphatidylinositol signaling system (Figure 5C). M12 was strongly associated with nodulation, peptidyl-serine modification, and protein phosphorylation processes, with circadian rhythm as the only identified KEGG-enriched pathway (Figure 5D). The absence of an enriched pathway related to nodulation in M12 may be due to the lack of information in the KEGG database.

### 2.5 Stand-out processes and pathways in the plant responses to nodulation, water deficit or the combination of both conditions

Combining information from different data layers may lead to new biologically interpretable associations (van Dam et al., 2018). Here, we further investigate the DEGs’ more relevant WGCNA modules for protein-protein interaction (PPI)-enriched networks, from which novel biological knowledge can be derived. The online STRING tool, which integrates known and predicted associations between proteins, including both physical interactions and functional associations (Szklarczyk et al., 2021), was used to construct different PPI networks.

On the one hand, two particular subsets of DEGs obtained in comparison *i* (N+WR *vs.* N+WW) and *ii* (N+WR *vs.* NN+WR) were analyzed to seek potential interactions between them. These particular subsets of DEGs are highlighted in bold in Table 1, and the PPI networks are depicted in Figure 6.

**Figure 6.**
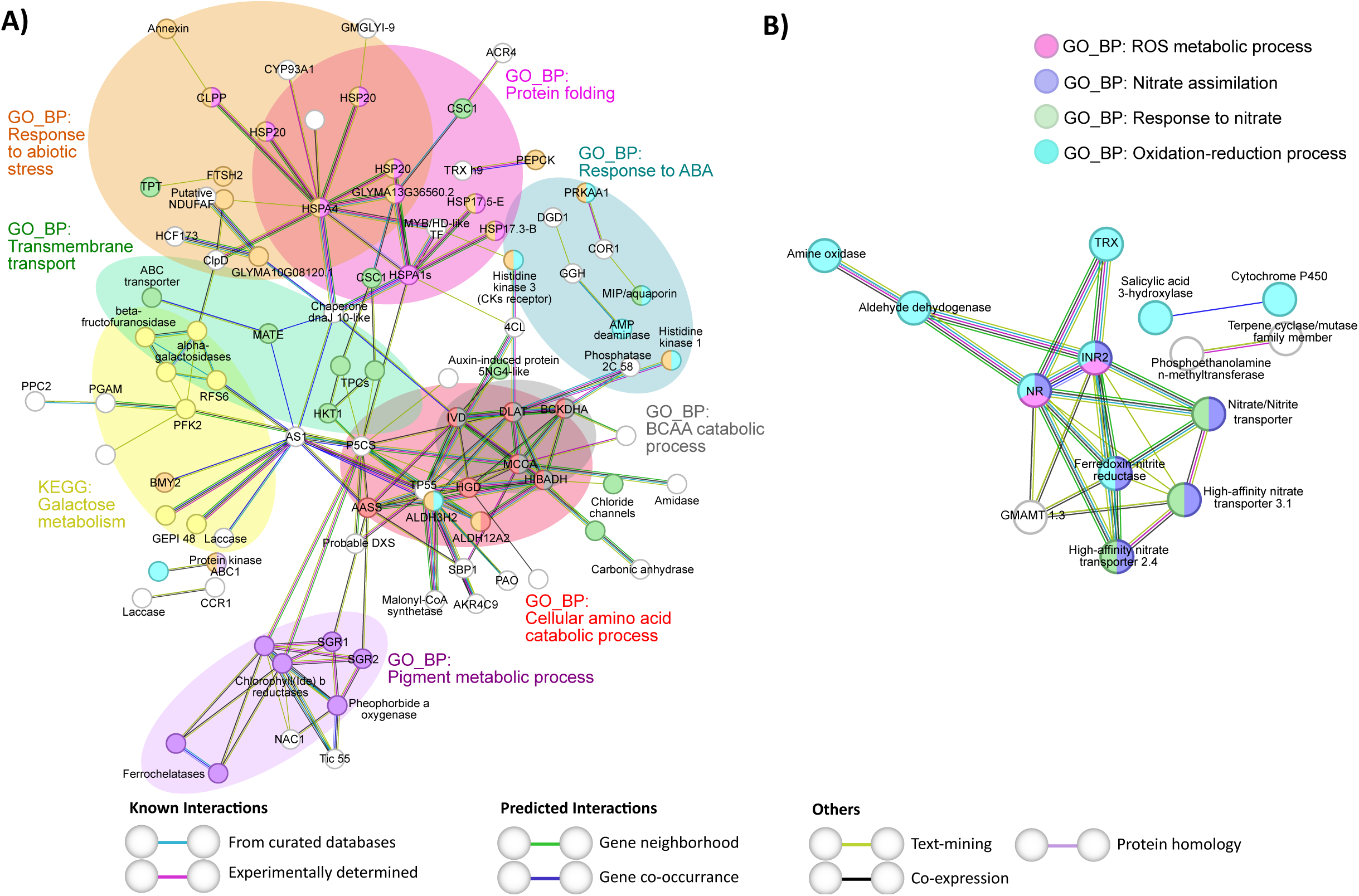
String analysis derived protein-protein interaction networks obtained from two subsets of differentially expressed genes discriminated by WGCNA modules at TOTAL+PAR level of comparisons *i* (A) and *ii* (B). **(A)** Network obtained from WGCNA M3 up-regulated genes in nodulated and water-restricted plants with respect to nodulated and well-watered plants (*i)* N+WR *vs.* N+WW contrast) (highlighted in bold in Table 1). **(B)** Network obtained from WGCNA M7 down-regulated genes in nodulated and water-restricted plants with respect to non-nodulated and water-restricted plants (*ii)* N+WR *vs.* NN+WR contrast) (highlighted in bold in Table 1). The network nodes represent proteins. Connecting lines denote protein-protein associations whose colors represent the type of interaction evidence as defined in the legend. The GO_BP and KEGG functional terms are also specified in the legends according to their colors. Disconnected nodes, i.e., proteins not showing any interaction in the network, were deleted.

In comparison *i*, the subset of DEGs selected comprised M3 up-regulated genes at TOTAL+PAR level (Figure 6A). The over-represented functional terms obtained in this PPI network depict the main processes that suffered changes due to up-regulation in nodulated plants subjected to water deficit, pinpointing them as candidates for further investigations. These processes can be mainly described by the following GO_BP terms and KEGG pathway: response to abiotic stress, protein folding, response to abscisic acid (ABA), transmembrane transport, pigment metabolic process, cellular amino acid catabolic process, branched-chain amino acids (BCAA) catabolic process, and galactose metabolism (Figure 6A). The first two BPs mentioned above shared many nodes (proteins), mainly heat shock proteins (HSPs) from different families. Among the proteins associated with the term transmembrane transport were ABC (ATP-binding cassette) transporters, calcium, chloride, and sodium channels, and MATE (Multidrug And Toxic Compound Extrusion) family transporters. The BCAA catabolism BP was entirely comprised within the cellular amino acid catabolism cluster; however, its identification highlights the importance of these small hydrocarbon-branched amino acids, which serve as precursors to secondary metabolites (DeKraker and Gershenzon, 2011) and whose breakdown process is relevant in plants subject to different environmental conditions such as long-term darkness (Peng et al., 2015) and water deficit (Pires et al., 2016). Nodes that exclusively belonged to the cellular amino acid catabolic process comprised proteins involved in the degradation of Phe, Tyr, Pro, and Lys. The proteins that were included in the galactose metabolism KEGG functional term are related to cell wall metabolism. The pigment metabolic process term mainly comprises proteins related to chlorophyll breakdown, such as stay green proteins, chlorophyll(Ide) b reductases, and pheophorbide a oxygenase (Figure 6A).

In comparison ii, the selected subset of DEGs corresponded to M7 down-regulated genes also at TOTAL+PAR level (Figure 6B). The PPI network of this particular subset of DEGs presented enriched terms related to nitrate metabolism (nitrate assimilation and response to nitrate), ROS metabolic process, and oxidation-reduction process (Figure 6B). In fact, in nodulated and water-restricted plants, the down-regulation of proteins related to nitrate uptake and reduction makes sense when compared to non-nodulated (nitrate-fed) plants.

On the other, the two other subsets of DEGs selected to seek potential interactions corresponded to comparisons *i* and *ii* common DEGs (Figure 7A) and to comparisons *ii* and *iv* common DEGs (Figure 7B). These two particular subsets of DEGs are highlighted in bold in Table 2. For comparisons *i* and *iii* common DEGs (related to water deficit responses, independently of the nodulation condition), the subset analyzed included M3 up-regulated genes at TOTAL+PAR level. For comparisons *ii* and *iv* common DEGs (related to nodulation, regardless of the hydric condition), the subset comprised M12 up-regulated genes, also at TOTAL+PAR level. The PPI network for common DEGs in comparisons *i* and *iii* revealed enrichment in four KEGG pathway functional terms: “glutathione (GSH) metabolism”, “plant hormone signal transduction”, “BCAA degradation”, and “galactose metabolism” (Figure 7A). Gray and yellow-outlined terms, BCAA degradation and galactose metabolism, were also present in the network depicting processes related to nodulated plants’ response to water deficit (Figure 6A). However, proteins associated with these terms varied between the two figures. The “GSH metabolism” term included glutathione peroxidase (GPx) and glutathione S-transferase (GSTs) enzymes, while “plant hormone signal transduction” involved protein phosphatases of the 2C family and an ABA-responsive element binding factor (bZIP71) (Figure 7A). The PPI network for DEGs related to the nodulation process (those common in comparisons *ii* and *iv*) showed enrichment in various processes, including “nodulation,” “interspecies interaction between organisms,” “inorganic ion transmembrane transport,” and “zeatin biosynthesis” (Figure 7B).

**Figure 7.**
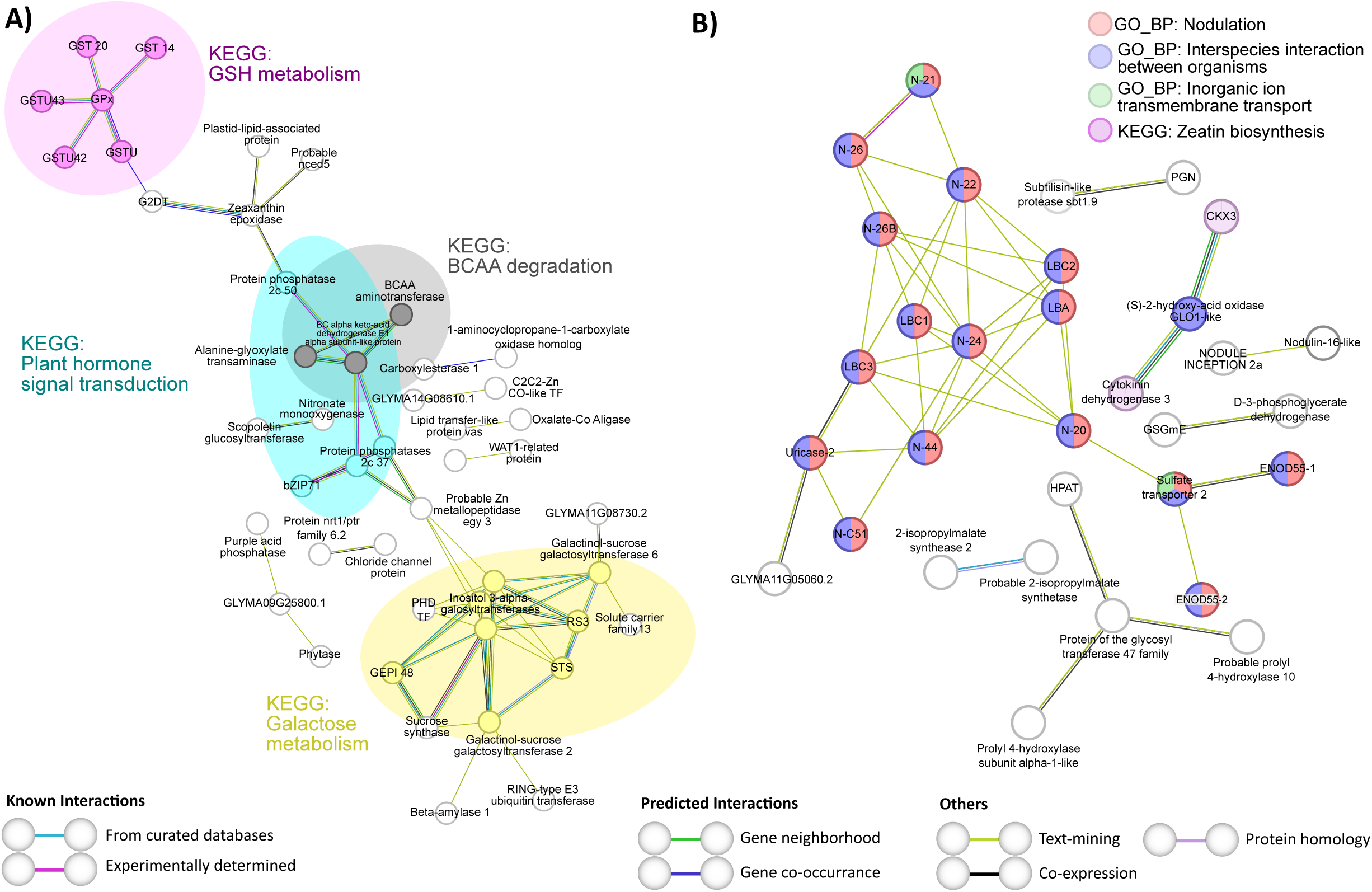
String analysis derived protein-protein interaction networks obtained from the differentially expressed genes common to contrasts *i*) and *iii*) (A) and common to contrasts *ii*) and *iv*) (B), discriminated by WGCNA modules at TOTAL+PAR level. **(A)** Network obtained from WGCNA M3 up-regulated genes (highlighted in bold in Table 2). (B) Network obtained from WGCNA M12 up-regulated genes (highlighted in bold in Table 2). The network nodes represent proteins. Connecting lines denote protein-protein associations whose colors represent the type of interaction evidence as specified in the legend. The GO_BP and KEGG functional terms are also specified in the legends according to their colors. Disconnected nodes, i.e., proteins not showing any interaction in the network, were deleted.

## 3. Discussion

Studies on how soybean plants respond to water-deficit stress have been mainly conducted on non-nodulated plants (Van Ha et al., 2015; Song et al., 2016; Wang et al., 2022), which in this work has been analyzed in comparison *iii* (NN+WR *vs*. NN+WW) (Figure 1C). However, all the major producer countries of soybean rely on the high symbiotic nitrogen fixation capacity of this plant (Maluk et al., 2023). Hence, evaluating the responses of non-nodulated legume plants to abiotic constraints is not the most relevant approximation if we aim to explore the crops’ response in the field. Moreover, evidence suggests that nodulated and non-nodulated plants respond differentially when subjected to water deficit (Borsani et al., 1999; Lodeiro et al., 2000; Staudinger et al., 2016; Liu et al., 2022), denoting even more, the relevance of dissecting how nodulated legume plants respond to water constraints. In addition to being different, the response of nodulated plants to water deficit is more complex in terms of the number of DEGs found (see Table S4, Table 1 and Figure 4A and C for comparisons *i* and *iii*). Actually, comparison *iii* (the one not including nodulation) had the lowest number of DEGs of the four comparisons analyzed (1881, 308, 263, and 523 DEGs for comparisons *i*, *ii*, *iii,* and *iv*, respectively; Table S4).

The differential response strategies—depending on the nodulation condition—of soybean plants to water deficit could be explained by gene expression regulation changes. In particular, translational control of gene expression allows cells to respond to a stimulus quickly, providing flexibility and adaptability to organisms (Hummel et al., 2009; Merchante et al., 2017; Sablok et al., 2017). Therefore, here, we analyzed the transcriptome and the translatome of the plants’ roots to identify genes subjected to translational control. ABA and ethylene are two crucial hormones that regulate plant responses to various stresses (Müller, 2021). Additionally, cytokinin and auxin play essential roles in controlling cell proliferation, differentiation, and processes like root nodule development in plants (Suzaki et al., 2013). Hence, the fact that we found DEGs mainly regulated at the translatome level in N+WR plants (with respect to N+WW and/or NN+WR plants) encoding various enzymes involved in ABA biosynthesis and signaling, such as zeaxanthin epoxidase and NDR1/HIN1-like protein 6, ethylene biosynthesis, like ACC synthase 3, which catalyzes the formation of ACC (a direct precursor of its biosynthesis), auxins—specifically IAA—inactivation, like IAA-amido synthetase GH3.10 and IAMT1, and cytokinin biosynthesis, such as IPT5, is very interesting, have not been addressed previously in our experimental conditions and is a focus point for future research regarding the fine-tuning of the molecular mechanisms governing the response of nodulated plants to water restriction (Figure 2, E, F; Table S2). Genes with mixed (transcriptional + translational) regulation are the most prevalent, so we performed the last part of our analysis (PPI network analysis; Figures 6 and 7) with the DEGs having the before-mentioned mixed regulation.

The analysis of our transcriptomic data through a pipeline that combines WGCNA+DEGs (Sánchez-Baizán et al., 2022) has proven effective for the identification of co-expressed gene modules (Figure 3) enriched with the DEGs obtained in the different comparisons. Interestingly, these DEGs co-localized specifically in some modules rather than being uniformly distributed throughout the WGCNA network (Figure 4; Table 1). This allows the association between module/s and processes underlying the plant responses being evidenced in each of the comparisons. An even more interesting aspect of this analysis that arises when the regulation status (up-or down-regulated) of the DEGs is analyzed is what is depicted in Figure 4 for comparisons *i* and *iv* (Figure 4A and D). It can be seen that many DEGs induced by nodulation and localized within M9, M10, M11, and M12 inverted their expression when water deficit was imposed on the nodulated plants. Notably, the biological processes represented in these modules were translation, DNA replication, regulation of transcription, protein ubiquitination, nodulation, and protein phosphorylation (Figure S2). This evidences once again the association between the DEGs’ enriched modules and the plant response mechanisms under evaluation and also that performing the differential expression analysis in the functional context of WGCNA facilitates the interpretation of complex scenarios as in the current study (Li et al., 2020b; Sferra et al., 2023). Likewise, this strategy was proper when we analyzed the DEGs common to comparisons *i* and *iii* and to *ii* and *iv*, i.e., the DEGs common to the plants’ responses to water deficit and nodulation, respectively (Table S5, Table 2). On the one hand, M3 and M7 can be assigned as the principal modules associated with the mechanisms that underlie the plants’ responses to water deficit independently of the nodulation condition (Table 2). Instead, M12 can be categorically assigned as the one module associated with the nodulation process, regardless of the hydric condition (Table 2).

To further investigate which molecular mechanisms were involved in the plants’ responses either to water deficit or nodulation, but mainly their combination, we performed GO_BP terms and KEGG pathways enrichment analysis (Figure 5) and PPI network analysis (Figures 6 and 7) for the most interesting modules and subsets of DEGs, respectively. M3 was identified as closely related to both the response of nodulated plants to water deficit—since most of the DEGs (91%) that localized on it correspond to contrast *i* DEGs (Table 1)—and, together with M7, to the plants’ responses to water deficit independently of the nodulation condition, considering that 85% of the common DEGs between contrasts *i* and *iii* localized within these two modules (Table 2). However, the stand-out processes and pathways in the before-mentioned plant responses differ both as to quantity and type of process or pathway, as can be seen in the PPI networks depicted in Figures 6A and 7A. As previously mentioned, terms related to “response to abiotic stress”, “protein folding”, “transmembrane transport”, “response to ABA”, “cellular amino acid catabolic process”, and “pigment catabolic process” were associated with the plant responses to nodulation + water deficit (Figure 6A). Instead, fewer terms were exclusively found to be associated with the plants’ response to water deficit: “GSH metabolism” and “hormone signal transduction” (2C protein phosphatases) (Figure 7A), which is in agreement with other studies (Van Ha et al., 2015; Lin et al., 2022; Ji et al., 2023). These findings support what has already been suggested (when considering the number of DEGs) about the response of nodulated plants to water deficit being unique and more complex if compare to that of non-nodulated plants.

Although two processes (“galactose metabolism” and “BCAA catabolism”) were common to both responses, the proteins associated with those terms in each case were different (Figures 6A and 7A). Proteins related to cell wall metabolism (included in the term “galactose metabolism”) in the nodulated plant’s responses to water deficit were UDP-glucose epimerases, which are involved in channeling UDP-D-galactose into cell wall polymers (Rösti et al., 2007), and alpha-galactosidase 3, which might have a role in cell wall loosening and expansion (Chuankhayan et al., 2023) (Figure 6A). Rather, proteins related to the term “galactose metabolism” in the plant’s responses to water deficit were inositol 3-alpha-galactosyltransferases (galactinol synthases), stachyose synthase (involved in the pathway that converts raffinose into stachyose), and galactinol-sucrose galactosyl transferases (also known as raffinose synthases). These enzymes have been shown to enhance drought tolerance in maize and Arabidopsis by raffinose synthesis or galactinol hydrolysis (Li et al., 2020a) (Figure 7A). The term “BCAA catabolism” in the response of nodulated plants to water deficit (Figure 6A) comprised mainly proteins associated with leucine, valine, and isovaleric acid catabolism, but also the branched-chain keto acid dehydrogenase E1 subunit alpha (BCKDHA) which is a component of the complex that catalyzes the second major step in the BCAAs catabolism. Interestingly, the DLAT (Dihydrolipoyl transacetylase) protein, a component of the pyruvate dehydrogenase complex, is also included in the “BCAA catabolic process”, confirming the connection that has been suggested between the degradation of these amino acids and the energetic status of the cells during drought conditions, considering they serve as alternative carbon sources as their degradation products are substrates of the tricarboxylic acid cycle (Pires et al., 2016; Zivanovic et al., 2020). Further, DLAT presented experimentally determined interactions with two ABA and abiotic stress-responsive histidine kinases: histidine kinase 1, for which a role as an osmosensor in water deficit conditions has been demonstrated in Arabidopsis (Wohlbach et al., 2008), and the cytokinin receptor histidine kinase 3. BCAAs have been suggested as important components of the ABA-mediated drought response (Zivanovic et al., 2020).

The relevance of identifying the transcriptomic and translatomic changes and related molecular mechanisms contributing to nodulated soybean plants’ responses to water deficit is clear. The experimental setup performed in this work allowed us to approach the above mentioned mechanisms through the up-regulated biological processes and metabolic pathways exclusively associated with nodulated and water-restricted plants. Moreover, the identification of the biological processes that, as a result of the nodulation process, were up-regulated but subsequently became down-regulated when the nodulated plants were subjected to water deficit shed light on the understanding of how nodulated plants respond to water deficit (Figure 8). Further comprehensive analysis of these processes could lead to identifying new sources of tolerance to drought in soybean.

**Figure 8.**
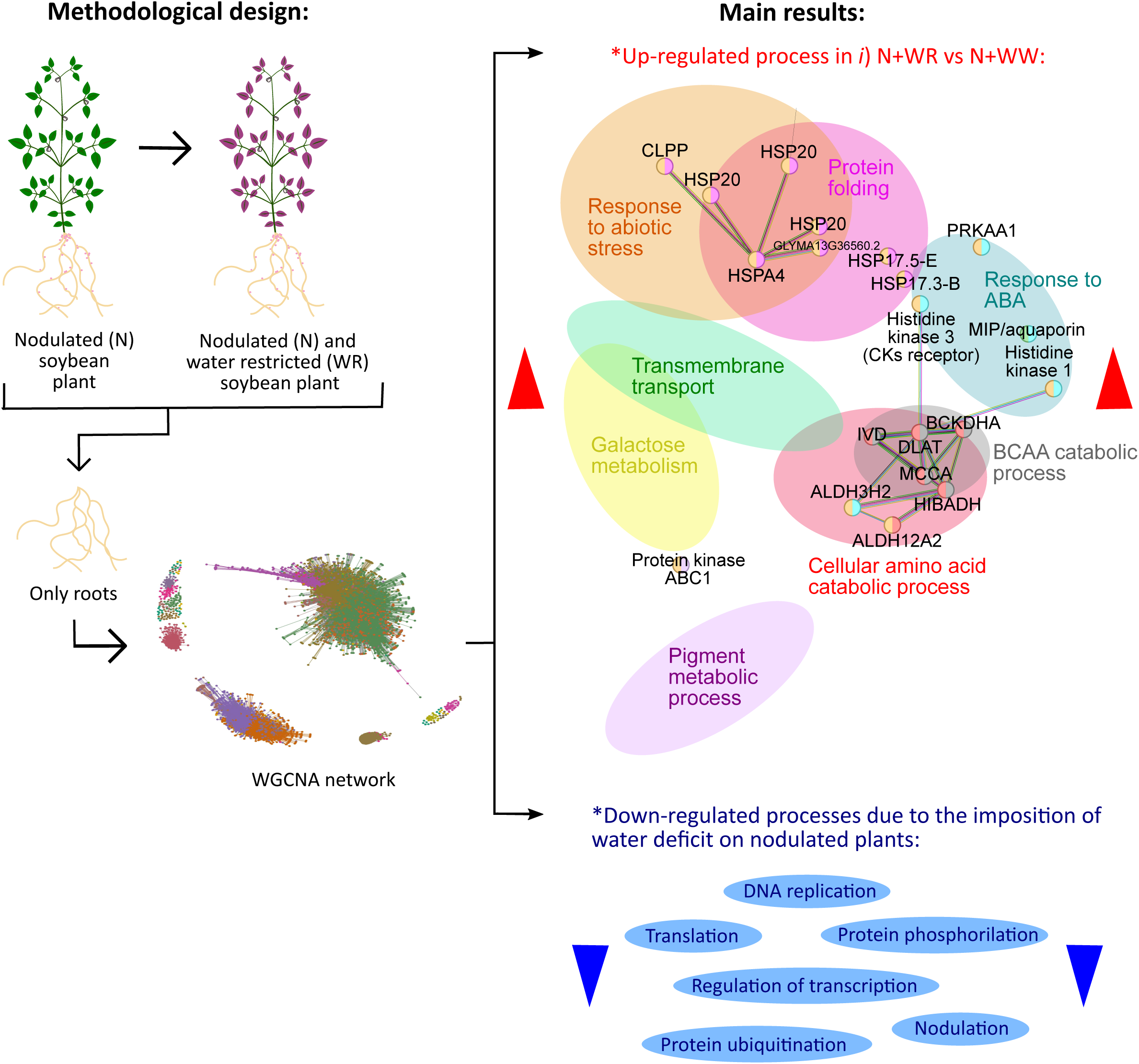
Schematic overview of the methodological design and main results of the work. Total and polysome-associated mRNA were extracted from the roots of N and WR soybean plants, along with their respective controls (NN and WW plants, not shown). The extracted mRNA underwent RNA-seq analysis, and subsequent weighted gene co-expression network analysis (WGCNA) was conducted. The figure depicts over-represented up-regulated functional processes linked to the response of N plants to WR (comparison *i*), as well as processes that were down-regulated due to the imposition of WR on N plants. N: nodulated; NN: non-nodulated; WR: water-restricted; WW: well-watered.

## 4. Materials and Methods

### 4.1 Plant Growth and Drought Assay

The assay was carried out with the commercial Don Mario 6.8i (DM) soybean [*Glycine max* (L.) Merr.] genotype as specified in (Sainz et al., 2022a). Briefly, plants were grown in a 0.5 L plastic bottle filled with a mix of sand:vermiculite (1:1) in a growth chamber under controlled temperature, photoperiod, and humidity conditions. The U1302 *Bradirhizobium elkanii* strain was used for the inoculated plants (Figure 1A). During the first 19 days after sowing (V2-V3 developmental stage), soybean seedlings were grown without water restriction keeping the substrate at field capacity with B&D-medium (Broughton and Dilworth, 1971) supplemented with KNO_3_ (0.5 mM and 5 mM final concentration for nodulated and non-nodulated plants, respectively). From day 20 after sowing, which corresponds to day 0 of the water deficit period, watering was withdrawn to the water-restricted (WR) plants. On the contrary, the well-watered (WW) plants were maintained without water restriction throughout the assay (Figure 1A). The substrate water content was measured daily by gravimetry (water gravimetric content) during the growth and water deficit period (Sainz et al., 2022a). The imposition of the water deficit was monitored through stomatal conductance (SC) measurements (Porometer Model SC-1, Decagon Device). Daily measurements were performed for all plants from day 0 until the end of the water deficit period, determined for each WR plant when the SC value was approximately 50% of the one obtained on day 0. At the end of the water deficit period of each WR plant, the roots were harvested and kept at -80°C until polysomal fraction purification. The roots of the WW plants were harvested together with the WR plants and kept at -80°C until polysomal fraction purification. At the moment of harvesting the nodulated plants, nodules were detached from the roots and both organs were kept at -80°C separately. Only the roots were analysed in this work.

The drought assay’s experimental design was a completely randomized one consisting of four combined treatments with five biological replicates each one, thus comprising a total of 20 pots. The experimental unit was one pot with one plant. The four combined treatments were nodulated (N) water-restricted (WR) plants (N+WR), nodulated well-watered (WW) plants (N+WW), non-nodulated (NN) water-restricted plants (NN+WR) and non-nodulated well-watered plants (NN+WW) (Figure 1B).

### 4.2 Polysomal Fraction Purification

The polysomal fraction purification by sucrose cushion centrifugation was performed according to (Smircich et al., 2015; Di Paolo et al., 2020; Sainz et al., 2022b) with some modifications. Two mL of packed volume of frozen pulverized roots were homogenized in 4 mL of polysome extraction buffer (see Sainz et al., 2022b for buffer composition) using mortar and pestle. Homogenates were maintained on ice with gentle shaking until all samples were processed and clarified by centrifugation at 16,000 g for 15 min. Next, the homogenate was filtered with cheesecloth and the centrifugation step was repeated. 500 µL of the supernatant was reserved for isolation of total RNA (TOTAL). Two mL of the remaining supernatant was loaded on 12% and 33.5% sucrose layers cushion in 13.2 mL tubes (Ultra-Clear, Beckman Coulter, United States, 344059) and centrifuge in a Beckman L-100K class S ultracentrifuge (W40 Ti swinging bucket rotor) at 4 °C for 2 h at 35,000 rpm. Refer to (Sainz et al., 2022b) for sucrose cushion stock solutions and layers preparation. After centrifugation, the polysomal fraction was recovered as a pellet and resuspended in 200 µL of polysome resuspension buffer (see Sainz et al., 2022b for buffer composition) pipetting up and down several times. The resuspended polysomal pellet was maintained for 30 min at 4°C and then regular RNA purification was performed to obtain the polysome-associated mRNA (PAR) fraction.

The confirmation of the presence of polysomes in the pellet obtained through sucrose cushion centrifugation was assessed through a continuous sucrose density gradient. The gradients were made from 15% and 50% sucrose solutions using a linear gradient maker and a peristaltic pump. Once cold, the centrifuge tubes containing the gradients were loaded with 200 µL of the resuspended polysomal pellet and centrifuged in the same conditions as the sucrose cushion centrifugation. The gradient fractions were analyzed using a fixed-wavelength (254 nm) detector (Cole-Parmer, United States, EW-42664-35).

### 4.3 RNA Extraction and Transcriptome Sequencing

TOTAL and PAR RNA fraction extraction was performed with TRizol LS reagent (Invitrogen, United States, 10296-028). The resuspended polysomal pellet and the extract reserved for isolation of total RNA were homogenized in 750 µL of TRizol, and the procedure shown in (Sainz et al., 2022b) was followed. After the resuspension of the RNA pellets, RNA concentration and integrity were measured using an Agilent 2100 bioanalyzer (Agilent Technologies, Inc., United States). Samples with a RIN (RNA integrity number) > 7.0 and > 1.0 µg were sent to Macrogen Inc. (South Korea) for library preparation and sequencing. TruSeq Stranded mRNA paired-end cDNA libraries were made and sequenced by the Illumina high-throughput sequencing platform. TOTAL and PAR samples from three biological replicates per combined treatment were sent for analysis.

### 4.4 Data analysis

#### 4.4.1 Sequencing read processing

Per sample sequencing quality was visually inspected using FastQC (https://www.bioinformatics.babraham.ac.uk/projects/fastqc/). Trimmomatic was used for adapter remotion and low-sequencing-quality bases trimming. Only those trimmed sequences longer than 80 bp, with overall quality higher than 30, were retained for further analysis. Gene expression, on a transcript level, was quantified using salmon quasi-mapping mode (v0.12.0, (Patro et al., 2017). Salmon default parameters were used, except the GC bias correction, which was enabled. The index for mapping was built from the most recent version of the reference *Glycine max* transcriptome, as retrieved from NCBI (GCF_000004515.6). Transcript read counts were then aggregated to gene level using the R package Tximport (v 1.2.0, Soneson et al., 2016). Descriptive data, as initial read counts, percentage of data retained after quality control, mapping rate, beside others, is presented in Table S1. The sequencing data is available in the NCBI Sequence Read Archive (SRA) under the accession number PRJNA868178.

#### 4.4.2 Differentially expressed genes (DEG) analysis

Initial descriptive analysis, as principal component analysis (PCA) of the samples and heatmap, were conducted using R base functions, and the R packages ggplot2 (Wickham, 2016) and pheatmap (Kolde, 2019). Statistical analysis of differential expression analysis was performed with DESeq2 (version 1.16.1, (Love et al., 2014), using the automated DESeq2 function. Differentially expressed genes (DEG) were defined as those with |log2FC| > 1 and Benjamini–Hochberg adjusted p-value (padj) < 0.05. Cluster analysis was performed using VennDiagramm (Chen, 2022), in order to identify DEG specifics for each contrast.

DEGs that co-localized with WGCNA modules of interest were subsequently used for Gene Ontology enrichment analysis, using topGO (Alexa and Rahnenfuhrer, 2023) and the weight01 method. Fisher’s exact tests were applied to the biological process (BP) category. GO terms with a FDR < 0.05 were considered significantly enriched. BP_GO enrichment plots, depicting the top-10 significant GO terms, were generated using the ’enrichment_barplot’ function of topGO. The gseKEGG function, as implemented in the clusterProfiler package (Wu et al., 2021) was used for KEGG pathway gene set enrichment analysis. The STRING tool (Szklarczyk et al., 2021; 11.5 version) was used to search for potential interactions between the DEGs.

#### 4.4.3 WGCNA analysis

A co-expression network analysis was performed using the WGCNA package (v1.71, (Langfelder and Horvath, 2008). For this analysis, DEseq2 (version 1.16.1, (Love et al., 2014); see below) normalized data was initially filtered, to remove genes with low coefficient of variation and/or low counts (< than 50 in more than 50% of the samples), as low-expressed features tend to reflect noise. Then, the pickSoftThreshold function in WGCNA was used to select the soft threshold power for the scale-free topology network. This way, the 1-step network construction and module detection function was applied, with power set to 17, maxBlockSize defined as the total number of genes, mergeCutHeight set to 0.20 and signed TOMType (as signed networks preserve the natural continuity of the correlation).

To determine whether the generated modules are associated with treatment groups, an Eigengene (hypothetical central gene) was calculated for each module, using the ‘moduleEigengenes’ function of WGCNA. A heatmap, depicting the relationship between module Eigengene and treatment groups, was generated using pheatmap (Kolde, 2019).

Network was generated using the R package network (Butts, 2008), with the edge adjacency set to 0.15. For visualization purposes, modules were projected in different colors. Alternatively, DEG genes (see *4.4.2*) were colored in the network, based on their regulation status (i.e. up/ down regulated). Gene connectivity was estimated using the ‘intramodularConnectivity’ function of WGCNA. Hub genes were defined as the top 10% genes of highest connectivity, following the recommendations of (Seo et al., 2009). Finally, transcription factor annotation was performed following Jin et al., 2017.

## Acknowledgments

We thank Joaquina Farías (Centro Universitario Regional NorEste-Universidad de la República) for helpful comments on the manuscript. We also thank the Sistema Nacional de Investigadores (ANII) (María Martha Sainz, Carla Valeria Filippi, Guillermo Eastman, Mariana Sotelo-Silveira, José Sotelo-Silveira, Omar Borsani).

## Author contributions

M.M.S., J.S.-S. and O.B. conceived the study; M.M.S., C.V.F., G.E., S. Z. and M.M.-M. performed the experiments; M.M.S., C.V.F., G.E., M.S.-S. and J.S.-S. analyzed the data; M.M.S. wrote the original draft except for the data analysis section of Materials and Methods; C.V.F. wrote that part. All authors have reviewed and edited the manuscript.

## Supplementary data

**Figure S1. Descriptive analysis of the RNA-seq data. (A)** Heatmap of all samples comprising three replicates of each combined treated sample (N+WR; NN+WW; N+WW; NN+WW) and RNA fractions (TOTAL and PAR). **(B)** Principal component analysis.

**Figure S2. Functional enrichment analysis of the co-expression modules most representatives of the differentially expressed genes (DEGs) in the different contrasts between treatments. (A)** M3. **(B)** M5. **(C)** M6. **(D)** M7. **(E)** M9. **(F)** M10. **(G)** M11. **(H)** M12. The top ten Gene Ontology Biological Process (GO-BP) terms are shown for each module (y-axis).

**Table S1. Sequencing general statistics.**

**Table S2. Fold change (FC) at the transcriptome (TOT) and translatome (PAR) level in each of the four comparisons (*i*, *ii*, *iii* and *iv*) among the combined treatments analyzed.** The status (up-or down-regulated) of the genes is shown.

**Table S3. Results of WGCNA and differential expression analysis.** The numbers depicted in columns F and G correspond to the Venn diagrams intersections (indicating contrast (i, ii, iii, iv) and regulation level (TOTAL, PAR, TOTAL+PAR) shown in the Figure in the next tab.

**Table S4. Differentially expressed gene (DEG) analysis coupled with weighted gene co-expression network analysis (WGCNA).** All 25 modules (M) are depicted. The four different contrasts (*i*, *ii*, *iii* and *iv*) in which the DEG condition was achieved are shown, as well as the status (up- or down-regulated) and the regulation level (TOTAL, PAR, or TOTAL+PAR).

**Table S5. Differentially expressed genes (DEGs) common to contrasts *i*) and *iii*) and contrasts *ii*) and *iv*) at TOTAL, PAR, or TOTAL+PAR level, coupled with weighted gene co-expression network analysis (WGCNA).** All 25 modules (M) are depicted, as well as their status (up- or down-regulated).

